# Multivariate analysis of EEG activity indexes contingent and non-contingent attentional capture

**DOI:** 10.1101/734004

**Authors:** Jaap Munneke, Johannes Fahrenfort, David Sutterer, Jan Theeuwes, Edward Awh

## Abstract

It is well known that salient yet irrelevant singleton can capture attention, even when this is inconsistent with the current goals of the observer (Theeuwes, 1992; 2010). Others however have claimed that capture is critically contingent on the goals of the observer: Capture is strongly modulated (or even eliminated) when the irrelevant singleton does not match the target-defining properties (Folk, Remington, & Johnston, 1992). There has been a long-standing debate on whether attentional capture can be explained by goal-driven and/or stimulus-driven accounts. Here, we shed further light on this phenomenon by using EEG activity (raw EEG and alpha power) to provide a time-resolved index of attentional orienting. Participants searched for a target defined by a pre-specified color. The search display was preceded by a singleton cue that either matched the color of the upcoming target (contingent cues), or that appeared in an irrelevant color (non-contingent cues). Multivariate analysis of raw EEG and alpha power revealed preferential tuning to the location of both contingent and non-contingent cues, with a stronger bias towards contingent than non-contingent cues. The time course of these effects, however, depended on the neural signal. Raw EEG data revealed attentional orienting towards the cue early on in the trial (>156 ms), while alpha power revealed sustained spatial selection in the cued locations at a later moment in the trial (>250 ms). Moreover, while raw EEG showed stronger capture by contingent cues during this early time window, the advantage for contingent cues arose during a later time window in alpha band activity. Thus, our findings suggest that raw EEG activity and alpha-band power tap into distinct neural processes that index movements of covert spatial attention. Both signals provide clear neural evidence that both contingent and non-contingent cues can capture attention, and that this process is robustly shaped by the target-defining properties in the current block of trials.

## Introduction

Two opposing views of attentional capture can be distinguished, based on the degree to which the internal goals of the observer influence this process. Some have argued for a fully stimulus-driven, bottom-up account of attentional capture in which attention is automatically and involuntarily allocated to the location of a salient stimulus, such as an abrupt onset (Schreij, Owens, & Theeuwes, 2008; Schreij, Theeuwes, & Olivers, 2010) or a stimulus with unique visual features such as its color or luminance (Kim & Cave, 1999; Theeuwes, 1991, 1992, 2004), which makes the stimulus “pop-out” from its surrounding elements. By contrast, others have argued that attentional capture is contingent on an observer’s current target template. This phenomenon is known as feature-based contingent capture and refers to the observation that attention is automatically captured by task-irrelevant stimuli that share vital visual features with the target stimulus (Bacon & Egeth, 1994; Egeth & Yantis, 1997; Folk & Remington, 1998; Folk et al., 1992; Leber & Egeth, 2006; Wolfe, Butcher, Lee, & Hyle, 2003).

Contingent capture was initially observed in a study by Folk and colleagues (1992), in which participants were instructed to detect a target that was either defined (in separate blocks of trials) based on its color (e.g. a red character among white characters) or by its presentation as an abrupt onset. Shortly (150 ms) before presenting the target display a brief cue display was presented consisting of a color or an abrupt onset stimulus, presented at one of four possible target locations. The location of the cue was task-irrelevant and did not predict the subsequent target location. The critical observation was that *only* the cue that shared critical features with the target stimulus captured attention, resulting in a strong validity effect (i.e. faster response times when the cue correctly indicated the target location, as compared to when cue and target were presented at different spatial locations). Crucially, when participants were searching for a target defined by an abrupt onset, a red cue amongst white distractors did not show evidence of attentional capture, whereas this cue did capture attention when participants were searching for a red target. This finding was taken as evidence that attentional capture is not merely bottom-up but that feature-based attentional mechanisms are instrumental in this process as well.

The finding that automatic capture is conditional on an observer’s target template is in stark contrast with the notion that attentional capture is a purely bottom-up, stimulus-driven phenomenon. To further investigate the nature of the contingent capture, Belopolsky and colleagues (2010) adapted the contingent capture paradigm used by Folk and colleagues (1992) such that the target defining feature was not kept constant over a number of trials, but rather provided the participants with a target template on a trial-by-trial basis by using an instructive cue. The results of this experiment showed strong attentional capture by relevant (matching the observer’s top-down set) as well as irrelevant (not matching the observer’s top-down set) distractors, providing evidence that contingent capture is ultimately bottom-up in nature and that the results by Folk and colleagues could be explained by bottom-up inter-trial priming.

Contrary to the behavioral evidence in support of a bottom-up account of attentional capture, recent EEG work has provided convincing evidence that attentional capture is contingent on the observer’s feature-based top-down set by measuring the N2pc. This is an event-related potential (ERP) component that indexes the side of space where a selected item appears. The N2pc is a negative deflection that onsets approximately 200 ms after stimulus onset and is observed over the parieto-occipital scalp sites contralateral to selected visual stimuli. Eimer and Kiss (2008) showed that the N2pc was elicited in response to a salient and contingent (i.e. target matching) cue, but only when the target was presented among distractors. The N2pc would not be observed in response to a salient contingent cue when the target was presented in isolation. Further evidence that the N2pc reflects feature-based attentional processes was provided in a recent study by Grubert and colleagues (Grubert, Fahrenfort, Olivers, & Eimer, 2017). They showed that non-salient stimuli elicited an N2pc component, but only when these stimuli shared critical features with the target *and* the observer was actively searching for this target. However, when the target was already found and the top-down set was no longer active, the same non-salient stimuli did not generate an N2pc. The findings observed by Grubert and colleagues were taken as evidence that the N2pc indeed reflects feature-based attentional processes, and the absence of an N2pc when the observer is not actively searching for a target cannot be explained in terms of bottom-up attention (but see Hickey et al., 2006, for a bottom-up interpretation of the N2pc).

Thus, EEG evidence has suggested that an observer’s target template influences attentional allocation by showing a tight link between the N2pc and feature-based attentional capture. However, one of the shortcomings of using the N2pc is that it is a transient neural response that occurs approximately 200 ms post stimulus onset, so it does not allow sustained tracking of covert attention during subsequent points in time. In addition, the N2pc as a measure of attentional allocation lacks spatial specificity (Fahrenfort, Grubert, Olivers, & Eimer, 2017). The N2pc is measured as a difference in electrical potential between two electrodes placed over ipsi- and contralateral cortical regions (relative to a visual stimulus) and as such does not provide information concerning the attended location beyond hemispheric differentiation. These shortcomings, as well as those imposed by purely behavioural research, may impede distinguishing between a bottom-up or feature-based nature of attentional capture. As such, the main goal of the current study was to investigate the mechanisms underlying attentional capture using a temporally resolved method that tracks attention throughout an entire capture trial. As the data will show, this provides new insights regarding early and late neural components that index attentional capture, as well as the impact adopting a target template has on this process.

To obtain a high-resolution spatiotemporal profile of attentional capture, we tracked the locus of covert attention using the scalp distribution of both raw EEG (Fahrenfort, Grubert, Olivers, & Eimer, 2017) and alpha power (Foster, Bsales, Jaffe, & Awh, 2017; Foster, Sutterer, Serences, Vogel, & Awh, 2016; Foster, Sutterer, et al., 2017). Recent work has shown that both signals provide time-resolved and precise tracking of currently attended locations, making this an ideal approach for understanding temporal dynamics of attentional capture. The current study uses this methodology to characterize the spatiotemporal properties of attentional capture by investigating how contingent and non-contingent capture develop over time. We note that phase-locked changes in the raw EEG distributed signal were investigated, whereas non-phase-locked changes were studied in the induced alpha-band power distribution (see Methods). To our knowledge, both neurophysiological correlates of attention (i.e., raw EEG and time-frequency information in the alpha-band range) have not been combined in a single study. Although both neural signals track covert attention and distinguish between contingent and non-contingent cues, our results will show the divergent time course of these effects, indicating that these signals may tap into distinct aspects of spatial selection.

## Methods

### Participants

We tested 33 participants (22 females; mean age ± SD = 22.48 ± 3.21 years) with normal or corrected-to-normal vision. All participants gave written informed consent prior to the start of the experiment. All participants were recruited from the student community of the University of Oregon, USA (25), and the student population of Bilkent University, Turkey (8). Participants received a monetary reward or course credits for completing the experiment. The experimental procedures of this and all subsequent experiments were approved by the ethical committees of the University of Oregon and Bilkent University, and in accordance with the Declaration of Helsinki.

Out of the 33 tested participants, four had to be discarded for either showing poor behavioral performance (accuracy around chance: one participant) or technical issues during measurement (three participants). All reported analyses below are based on data from the remaining 29 subjects.

### Stimuli and Procedure

The experiment was conducted at the University of Oregon, USA and Bilkent University, Turkey, with near-identical procedures between the testing locations^1^. Participants were seated in a dimly lit room, at a viewing distance of 65 cm from a 22” CRT computer monitor. Prior to the start of the experiment, a 20-channel EEG electrode cap was fitted on the scalp of the participants and attached to an SA Instrumentation amplifier located in a Faraday cage.

Figure 1 shows the time course of a typical experimental trial. Participants started the trial by fixating on a centrally presented gray fixation cross (0.3° × 0.3°). After 500 ms, the fixation cross turned black for 100 ms, as a general indication that the critical part of the trial had started and that participants should refrain from making eye movements until the end of the trial. The fixation cross turned back to gray for a random period between 800 to 1200 ms (in increments of 100 ms), after which the cue screen was presented. The cue screen consisted of eight circles (2.5° radius) presented in a circular array around fixation with a radius of 5.0 degrees of visual angle. On each trial, one of the circles was presented as a solid red or green disc (luminance 30 cd/m^2^), functioning as a non-predictive cue. After 50 ms, all circles were removed from the screen for either 100 or 600 ms during which only the fixation cross remained present. Following this fixation-only interval the search array was presented for 50 ms, consisting of eight circles, each having a small opening on the right or the left side. On each trial, the target circle was presented in a predefined color that remained constant throughout the experiment (red and green, counterbalanced over participants). Participants were instructed to give a speeded response indicating on which side the circle had a small opening.

**Figure 1.**
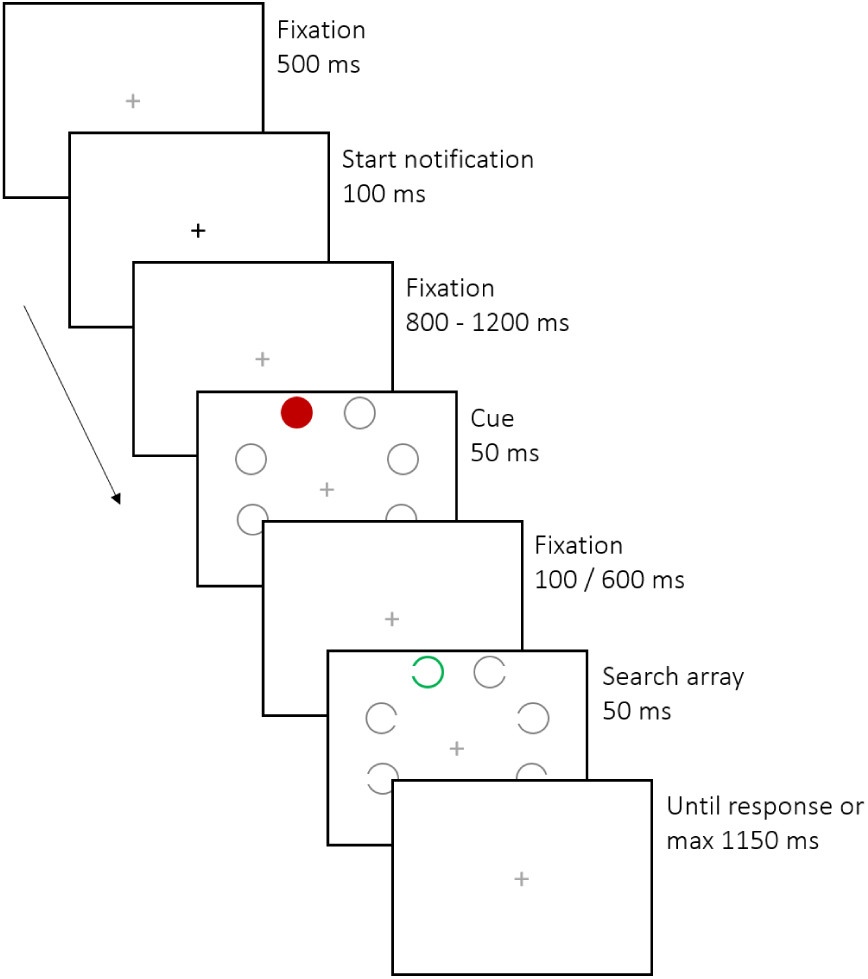
Time course of a typical experimental trial. In this particular trial, the cue is non-contingent (the colors of the cue and target do not match) and valid (cue and target are presented at the same location).

Short and long inter-stimulus intervals (Stimulus Onset Asynchrony: SOA) were used such that trials with a short SOAs (cue onset – target onset: 150 ms) reflected classic studies on contingent capture in order to illustrate any behavioral effects. Trials with a long SOA (650 ms) were used to investigate how contingent and non-contingent cues influenced automatic effects of capture, as well as spatial biases that were sustained for longer periods of time (up to 1000 ms after cue onset). Trials with short and long SOAs were randomly intermixed in each experimental block.

Cues were either contingent (having the same color as the target) or non-contingent (having a different color as the target). An equal number of contingent and non-contingent cues were used by counterbalancing the number of red and green cues and randomizing their order within the experiment. Furthermore, cue and target location were fully counterbalanced and presented equally often at each of the eight locations in the visual field, resulting in a cue (location) validity of 12.5%. The experiment consisted of 1280 (71.4%) trials with long SOAs and 512 (28.6%) trials with short SOAs. Finally, the eight cue and target locations were not fixed, but could be presented anywhere on the radius around fixation, with the limitation that the inter-stimulus distance remained constant at 4.14° degrees (i.e., the whole display could rotate, but the individual circles were always presented equidistant at a 45° angle between the center of the screen and two adjacent stimuli). Despite this random placement, the circle on which the stimuli were presented was divided in eight equally large segments, with each segment representing one location, despite the precise position of a stimulus within this segment. Therefore, we will refer to these segments as locations from here on. The entire session, including EEG preparation took approximately 2.5 hours to complete.

### EEG Recording and Preprocessing

A 20-channel electro-cap (Electro-Cap international) was used to record EEG from the following electrodes: F3, FZ, F4, T3, C3, CZ, C4, T4, P3, PZ, P4, PO3, PO4, PO7, PO8, POZ, T5, T6, O1, and O2. The left mastoid was used as an online reference and all data was re-referenced offline to the average of all electrodes. HEOG was obtained by placing two electrodes at the outer canthi of both eyes, enabling the measurements of horizontal eye movements. VEOG and blinks were measured by placing an electrode above and below the left eye^2^. All incoming signals (EEG and EOG) were amplified and filtered with a bandpass filter of 0.01– 80 Hz. Subsequently, all signals were online resampled at 250 Hz in LabVIEW 6.1 running on an external PC. Impedances were kept below 5 kΩ throughout the experiment.

Next, the EEG data was segmented in 2s epochs around cue onset (−500:1500). The contribution of eye blinks to the EEG signal was removed from the epoched data using an independent component analysis (ICA), by removing components that showed clear blink-related activity. Next, all ICA corrected epochs that contained data from trials with behaviorally incorrect responses were removed. The remaining epochs were checked for eye movements made in the time window 0-650 ms (i.e., from cue onset until target onset). Eye movements were detected by moving a 50 ms window over the preprocessed EEG data (in the 0 – 650 ms period for which we measure attentional effects on the probe) in steps of 50 ms within the HEOG or VEOG channels. Amplitude changes of 25 µv within that window were flagged as eye-movements and any trial containing such artifacts were subsequently deleted from the data set (2.95 %). Finally, epochs containing muscle artifacts were removed from the data by calculating the z-value of the power values in the EEG signal, for frequencies above 110 Hz (up to 125 Hz). Trials that contained *z*-score outliers more than 3 standard deviations away from the absolute value of the minimum negative *z*-score were marked as containing a muscle artifact and were removed from the data set (3.99 %). All preprocessing steps were conducted using EEGLAB (Delorme & Makeig, 2004) and the Amsterdam Decoding and Modeling toolbox (Fahrenfort, van Driel, van Gaal, & Olivers, 2018).

### EEG Analysis

In order to use alpha-band power in our analyses, FieldTrip (Oostenveld, Fries, Maris, & Schoffelen, 2010) was used to decompose the raw EEG signal into frequency-specific power spectra. Frequency-specific power spectra were based on a Fast Fourier Transform (FFT) approach using a fixed (i.e., independent of frequency band) 500 ms moving Hanning window (step size = 8 ms), resulting in a frequency resolution of 2Hz (1/0.5 sec). As such, the FFT analysis resulted in the time-frequency bins for all even frequencies ranging from 2 to 30 Hz (i.e., 2 Hz, 4 Hz, 6 Hz … 30 Hz). We calculated changes in induced (non-phase locked) power for each frequency and time point. The study specifically focused on changes in induced power to ensure that stimulus specific phase-locked signals found in the raw EEG were not present in this signal, thus measuring qualitatively different task-related signals than those encoded in the phase-locked EEG. Induced power was computed by subtracting the condition-specific average evoked response (ERP) waveform from each trial of that condition (cue-position and cue-type) prior to computing the signal’s power. This method effectively subtracts out the phase-locked part of the signal from every single trial, leaving only stimulus induced power fluctuations of signals that are plausibly already ongoing (hence non-phase locked) when the stimulus appeared. Induced power signals that are computed by subtracting out ERPs prior to time-frequency decomposition have (by algorithmic logic), a different ontology from signals contained in the phase locked, raw EEG. As such, any difference in classification performance between raw EEG and power-based analyses likely reflect expressions from distinct cortical mechanisms. Indeed, evoked (phase-locked) components such as the N2pc are not present in the induced signal, whereas induced signals are often thought to reflect endogenous process that are modulated, but not initiated by external stimulation or task instruction (e.g. David, Kilner, & Friston, 2006; Hosseini, Bell, Wang, & Simpson, 2015). The subtraction procedure that was used to obtain induced signals was applied separately to training and testing data in each fold, computing condition-specific ERPs for every training and every testing set, as to prevent the ERP subtraction method from inadvertently introducing commonalities into the entire dataset that could drive above-chance decoding across training and testing. Although it would strictly be sufficient to apply this procedure to the training data only, we chose to apply it to the testing set too, eradicating any remnants of phase-locked activity. However, because the test data did not have enough trials to allow the computation of a sufficiently clean ERP, we fitted a spline through test-set ERPs to remove high-frequency noise prior to subtracting them from the single trials in the testing set.

All analyses were multivariate, either applied to the raw EEG or to the time-frequency decomposed induced signal of the EEG. We first used backward decoding models (BDM) to infer whether we could predict the cue location based on the distributed EEG patterns. Next, we applied forward modeling techniques (FEM) to determine whether the underlying multivariate signal contained continuous tuning characteristics, and whether these differed for contingent and non-contingent. All BDM and FEM analyses were conducted using the Amsterdam Decoding and Modeling Toolbox (ADAM, Amsterdam, the Netherlands, Fahrenfort et al., 2018), which uses EEGLAB as input format and internally uses FieldTrip to perform time-frequency analysis.

#### Backward Decoding Model

A backward decoding model was used to predict at which of the eight possible locations the cue was presented, based on the distribution of EEG activity (raw EEG and time-frequency power distributions (alpha)). The underlying logic of this analysis is that if a trained classifier can predict with above-chance classification accuracy where the cue was presented, then it follows that location specific information is present in the distributed EEG patterns.

In order to conduct the BDM analyses on the data, a number of steps were taken to ensure the validity of the model. First, the trial order was randomized offline for every subject, to prevent order effects from affecting classifier performance in any way. Next, each subject’s individual dataset was analyzed using a 10-fold cross-validation training-testing scheme. In this scheme, the data was segmented into 10 equally sized folds (each fold containing a near-equal number of trials, with equal distributions of the eight cue positions across the folds). A linear discriminant classifier was trained on 90% of the data (9 of the 10 folds) learning to discriminate between the different stimulus classes (i.e., the 8 possible cue locations) separately for each of the two cue types. The validity of the trained classifier was tested on the left-out 10% of the data (the remaining fold); a procedure which was repeated ten times, such that all data was tested once without ever using the same data for training and for testing.

Separate decoding analyses were conducted using 1) the distributed amplitudes of the raw EEG signal over each electrode and time point and 2) the induced power of decomposed frequency information at each electrode and time point. Using 20 electrodes thus resulted in 20 features for 8 stimulus classes (8 cue locations), classified in two conditions (contingent and non-contingent cues). Rather than using the average proportion of correctly classified stimulus categories as a performance measure (e.g. Fahrenfort, Leeuwen, Olivers, & Hogendoorn, 2017), the BDM analyses used a slightly more sensitive performance measure by assessing the area under the curve (AUC; Hand & Till, 2001) of a Receiver-Operator Characteristic (ROC) that plots the cumulative probabilities that the classifier assigns to instances coming from the same class (i.e. the correct cue location) against the cumulative probabilities that the instance is classified as being from a different class (i.e. one of incorrect cue locations). As more than two classes (i.e. the cue location) were used, AUC was defined as the average AUC of all pairwise comparisons between classes. Using AUC as a performance measure is more sensitive than using decoding accuracy, as it uses single trial confidence scores (i.e. the distances from the decision boundary) to compute performance, rather than averaging the performance on a set of binary classifier decisions. AUC typically runs from 0.5 (chance performance) to 1 (perfect performance).

#### Forward Encoding Models (FEMs)

Compared to BDMs, forward encoding models (FEMs; Brouwer & Heeger, 2009) take the opposite approach by establishing the continuous relationship between a stimulus parameter (cue position in this case) and multivariate neural patterns. This relationship is expressed in a single so-called Channel Tuning Function (CTF, loosely reminiscent of tuning properties of single neurons), which together with regression weights obtained during model creation allows one to reconstruct neural patterns for stimulus parameters that were never used to create the model (Fahrenfort et al., 2017; Brouwer & Heeger, 2009). Hence, the term ‘forward model’ reflects the fact that one can go from the stimulus parameter space to predict neural activity (and vice versa).

During FEM model fitting, a similar 10-fold procedure is used as during backward decoding, but using a different procedure. First, a basis set is created for each of eight hypothetical channels reflecting cue position, and which describe the assumed (hypothetical) relationship between neural activity and the eight cue positions on the screen. The nomenclature “channels” here should not be confused with MEG or EEG sensors, EEG sensors are referred to as electrodes in the current manuscript. We used a Gaussian shaped basis set, which was created using a standard Gaussian function with an amplitude of 1 and a sigma of 1. Next, linear regression-based weight estimation for each of the hypothetical location channels was performed separately for each of the 20 features, specifying the one-to-one and invertible relationship between a particular cue position and the distributed multivariate neural response in the training set. Next, these weights were multiplied with trials in the testing set to produce the estimated channel responses for each trial in the testing set. This procedure was repeated for each of the 10 testing folds so that channel responses were derived once for each trial in all folds. Subsequently, the trial-based channel responses were averaged across trials in the testing set, separately for trials reflecting each of the eight different cue locations. The averaged channel responses in combination with the derived channel weights describe the validated and inversible relationship between attended cue location and the multivariate EEG response. In a final step, the eight estimated channel responses were aligned to a common center such that all eight channel responses were similarly centered. This step was conducted separately for each of the two cue conditions and was repeated for each time point, resulting in a CTF-over-time. The full procedure has been described at length in a number of other papers, both in mathematical terms (Brouwer & Heeger, 2009; Foster et al., 2016; Garcia, Srinivasan, & Serences, 2013; Samaha, Sprague, & Postle, 2016) as well as using more verbal and visual descriptions (Fahrenfort et al., 2017; Foster, Sutterer, et al., 2017). As in the BDM analyses, separate FEM analyses were conducted using the distributed raw EEG signal and the induced power spectra as input.

## Results

### Behavioral Results

#### Reaction Times

Only trials with correct responses were used in the reaction time analyses (5.23% discarded). Furthermore, for all analyses, trials with response times shorter than 200 ms as well as reaction times that were two standard deviations above the subject’s conditional means (4.11%) were removed. To investigate the effect of contingent and non-contingent cues on attentional allocation to the target presented in the search array, we first calculated the mean reaction times per condition for trials with a short SOA (150 ms). The time course in this condition best reflects the classic studies on contingent capture and allows us to draw conclusions about cue-induced attentional effects on target selection. Figure 2 shows the mean reaction times and accuracy scores for valid and invalid cues, separately for contingent and non-contingent trials. A repeated measures ANOVA on reaction times with these factors showed a main effect of validity, indicating that participants were faster on trials in which the location of the cue matched the location of the target, compared to when cue and target were presented at different locations (*F*(1,28) = 22.182, *p* < .001, *η*_*p*_^2^ = .442). No main effect of contingency was observed (*F*<1), but as expected, we did observe a clear interaction between contingency and validity (*F*(1,28) = 6.889, *p* = .014, *η*_*p*_^2^ = .197), with post-hoc t-tests showing that the validity effect was larger on contingent trials (Δ 24 ms; *t*(28) = 5.211, *p* < .001) as compared to non-contingent trials (Δ 8 ms; *t*(28) = 1.836, *p* =.077).

**Figure 2.**
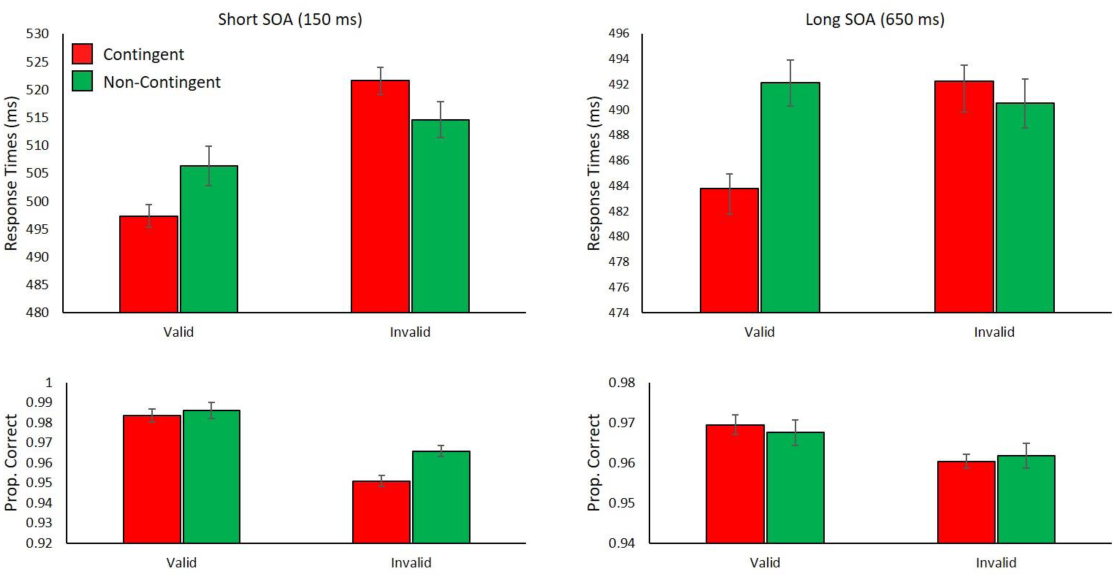
Mean reaction times and accuracy scores for the different experimental conditions. Separate plots are shown for trials with short and long SOAs. Error bars reflect the 95% confidence interval (Cousineau, 2005; Morey, 2008)

A similar analysis was conducted for all trials with a longer SOA (650 ms) in order to see if any of the cue-induced capture effects lingered such that it would influence the reaction times to targets presented with a larger temporal separation from the cue. No main effect of validity was observed (*F*(1,28) = 2.826, *p* =.104, *η*_*p*_^2^ = .092), suggesting that some of the attentional effects may have dissipated by the time the target was presented. Furthermore, a significant effect of contingency was observed (*F*(1,28) = 4.292, *p* =.048, *η* _*p*_^2^ = .133), with faster reaction times for contingent compared to non-contingent trials (see Figure 2 for means). Finally, similar to the analysis on the short SOA data, an interaction between contingency and validity (*F*(1,28) = 11.528, *p* = .002, *η* _*p*_^2^ = .292) was found. Post-hoc testing showed that this interaction was driven by the presence of a validity effect on contingent trials (Δ 8 ms; *t*(28) = 3.604, *p* =.001) that was completely absent on non-contingent trials (Δ −2 ms; *t*(28) = 0.607, *p* = .549).

#### Accuracy

Overall accuracy was relatively high: 94.60% correct. An ANOVA on the mean accuracy scores with contingency and validity as factors for the short SOA trials (150 ms) showed a main effect of validity, indicating that participants were more accurate on trials with validly cued targets as compared to trials with invalidly cued targets (*F*(1,28) = 33.795, *p* <.001, *η*_*p*_^2^ = .547; See Figure 2 – bottom panels for accuracy scores). A main effect of contingency was observed, indicating that participants responded less accurately on trials in which the target was preceded by a contingent compared to a non-contingent cue (*F*(1,28) = 8.976, *p* = .006, *η*_*p*_^2^ = .243). A significant interaction between contingency and validity was observed (*F*(1,28) = 4.212, *p* <.001, *η*_*p*_^2^ = .547), showing larger differences in accuracy between valid and invalid trials for contingent (3.27%; *t*(28) = 5.126, *p* < .001) as compared to non-contingent trials (2.0%; *t*(28) = 4.639, *p* <.001). A similar ANOVA on the trials with long SOAs (650 ms) showed only a significant main effect of validity, indicating that participants were overall more accurate on trials with validly cued targets as compared to invalidly cued targets (*F*(1,28) = 6.748, *p* = .015, *η*_*p*_^2^ = .194). No main effect of contingency (*F*(1,28) = 1.613, *p* = .215, *η*_*p*_^2^ = .054), nor an interaction between the two factors was observed (*F*(1,28) = 1.907, *p* = .178, *η*_*p*_^2^ = .064). Accuracy results need to be interpreted tentatively as the absence of certain effects may be masked or distorted by a ceiling effect due to the overall high accuracy of most of the subjects.

### EEG Results

#### Raw EEG analysis (BDM)

To first establish whether we could find early effects of attentional capture (i.e., the effects in the N2pc domain) and to gauge the extent to which these effects are shaped by cue contingency, we applied a backward decoding analysis to determine cue position using the raw EEG signal (similar to Fahrenfort et al., 2017b). Figure 3 shows classification accuracy (i.e. decoding accuracy as indexed by the ROC’s area under the curve) over time, indicating the extent to which the cue location could be predicted based on the multivariate raw EEG patterns. Classification accuracy (AUC) was tested against chance level (50%), separately for trials containing contingent (red) and non-contingent (green) cues. T-tests against chance were conducted for every time sample in the epoched data, correcting for multiple comparisons using 1000-iteration cluster-based permutation tests (see Maris & Oostenveld, 2007). As can be observed from Figure 3, both contingent (red) and non-contingent (green) cues yielded significant above-chance decoding performance (cluster-based *p* < .05, two-sided) emerging in the same time window as classical N2pc effects peaking between 200-250 ms after cue onset. To directly compare whether classification performance differed for contingent and non-contingent cues, classification performance for both cue types were tested against each other using paired samples t-tests. Again, these tests were conducted for each time point in the cue target interval, using cluster-based permutation testing to mitigate the multiple comparisons problem. Figure 3 shows the difference between classification performance for contingent and non-contingent trials (grey line / right axis; i.e., contingent – non-contingent) and clear significant differences for contingent compared to non-contingent cues can be observed in three consecutive temporal intervals (smallest cluster-based *p* < .001, one-sided), starting shortly after cue onset (156 ms after cue onset) and ending well before the targets appeared (420 ms after cue onset). Note that the peak difference of the difference wave is consistent with the time of maximum classification accuracy for both contingent and non-contingent cues.

**Figure 3.**
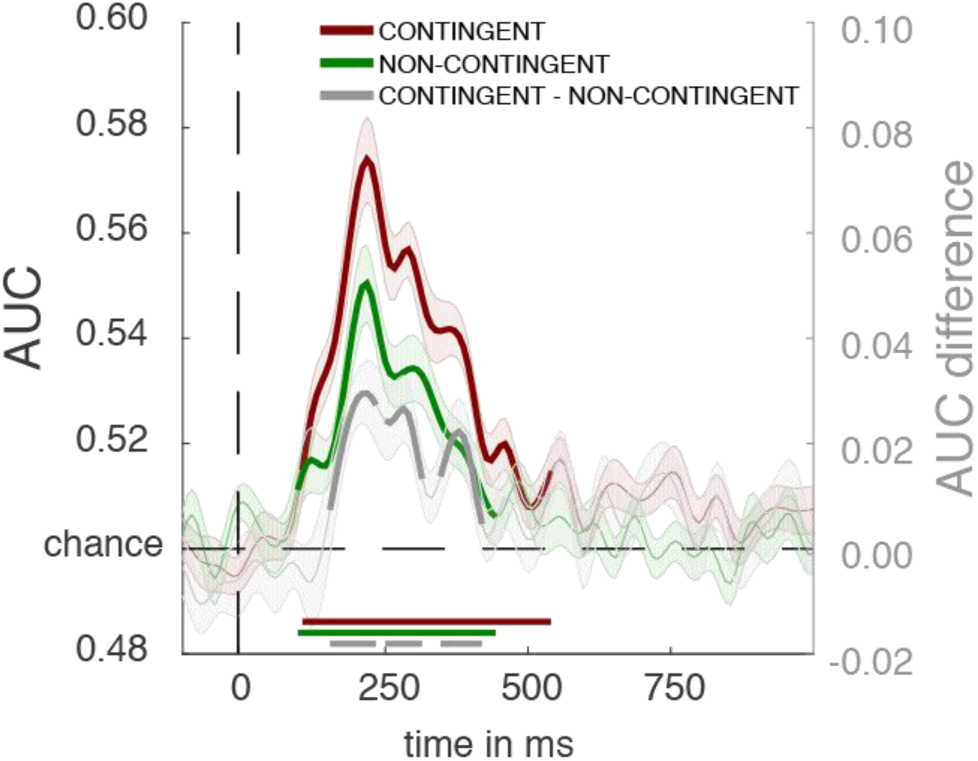
Classification accuracy using BDM, expressed as Area Under the Curve (AUC) for contingent (red) and non-contingent (green) trials as well as their difference (grey), based on the raw EEG signal. Colored bars on the x-axis show the intervals where classification performance is above chance (50.0%). Significant differences between contingent and non-contingent classification performance can be observed in multiple clusters ranging from 156 – 420 ms after cue onset (time point 0).

#### Time-Frequency analysis (BDM)

We applied a backward decoding analysis to the time-frequency data, aimed at investigating to what extent the multivariate distribution of the EEG’s power spectra could be used to predict at which location the contingent and non-contingent cues were presented. Past work has shown that this approach can track sustained orienting of covert attention, thereby providing an important complement to the analysis of the phase-locked EEG activity that appears to be most sensitive to attention effects occurring early after cue or target onset. Crucially, although separate multivariate analyses were conducted using the distribution of activity in the raw EEG on one hand and the distribution of power spectra amplitudes on the other hand, both measures were derived from the same data set and the same interval was tested for differential effects of the two cue types.

A first step in using distributed power spectra to decode the cue locations consisted of determining which neural oscillatory frequencies could be effectively used to decode this information. Therefore, a BDM analysis was conducted on all frequencies ranging from 2 – 30 Hz (in steps of 2 Hz; see methods), separately for trials with contingent and with non-contingent cues. Figure 4 shows the performance of the classifier for cue location, based on the induced (non-phase-locked, see methods) power spectra, over time. Note that the induced signal does not contain any of the phase-locked evoked responses that are present in the raw EEG data (see Methods for details). As expected, the highest decoding accuracy was observed in the alpha-band range for both contingent (Figure 4A) and non-contingent (Figure 4B) cues (Foster et al., 2016). As such, we used this frequency band (8-12 Hz) to further examine whether we could decode the location of the different cue types based on time-frequency information.

**Figure 4.**
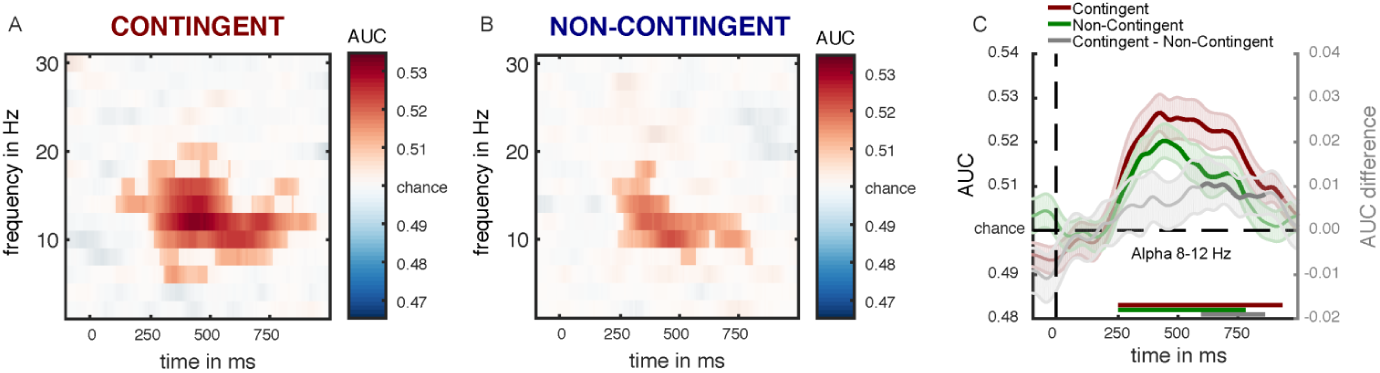
Classifier performance based on decomposed frequency power for frequencies ranging from 2 – 30 Hz (in steps of 2 Hz) for **A.** contingent and **B**. non-contingent (B) trials. Saturated values are cluster-corrected using cluster-based permutation testing (*p* < .05). **C.** Classifier accuracy over time based on 8-12Hz alpha-band power independently for contingent and non-contingent data. Green and red bars at the x-axis indicate the regions of above-chance decoding accuracy. The grey bar indicates the difference in decoding accuracy between contingent and non-contingent cues (scaled on the grey axis on the right). The difference between contingent and non-contingent is significant in an interval ranging from 596 – 860 ms after cue onset.

Figure 4c shows the above-chance classifier accuracy for contingent and non-contingent cues based on the distribution of alpha power over the scalp (red and green lines). Results showed that both contingent and non-contingent trials yielded above-chance classifier performance starting at approximately 250 ms post-cue and extending well past target onset (t_target_ = 650 ms). Crucially, the difference between contingent and non-contingent trials, as indicated by the grey line, was observed to be significant only in the later stages of the trial during an interval ranging from 596 ms to 860 ms post cue (cluster-based *p* = .036, one-sided).

Thus, decoding based on alpha power suggested a pattern of attentional engagement in which attention was rapidly allocated to the location of both contingent and non-contingent cues, with contingent cues yielding significantly higher decoding accuracy later in the trial (appearing shortly before target onset). When directly comparing the decoding accuracy based on alpha power for contingent and non-contingent cues, it appears that location tuning declined more quickly for non-contingent cues than for contingent cues. To further investigate whether a more specific model could bring out these differences between the two conditions we applied an FEM model to the raw EEG and the time-frequency data.

#### Raw EEG Analysis (FEM)

The early above-chance classifier performance for contingent and non-contingent cue locations was taken as an incentive to investigate whether a forward encoding model (FEM) could be used to create location-selective channel tuning functions (CTFs) that describe the continuous relationship between multivariate patterns of EEG activity and attended cue location, separately for contingent and non-contingent cues. As raw EEG is used as input for the forward model, we expect any differences between contingent and non-contingent to arise as early effects in the channel tuning response over time, reflecting early and automatic effects of attentional capture.

A forward encoding model that describes the relationship between the multivariate EEG patterns and cue location (separately for cue type) was constructed (see methods). Figure 5A shows how the CTFs for contingent and non-contingent cues develop in a 1000 ms time window following cue onset. Significance testing of the CTFs was conducted by testing the slopes of the CTF function against 0 for each time point, again corrected for multiple comparisons using cluster-based permutation testing (*p* < .05). The slope of each CTF was estimated using linear regression after collapsing across channels that were equidistant from the channel tuned to the cued location (i.e., channels −4 and 4, channels −3 and 3, etc. as reported in Figure 5). Significant periods in the developing CTFs are indicated by black bars at the bottom of the plots. As can be seen for both contingent and non-contingent cues, CTFs reach significance shortly after cue onset (approximately after 150 ms), dovetailing the observed time course of the BDM results using the raw EEG data.

**Figure 5.**
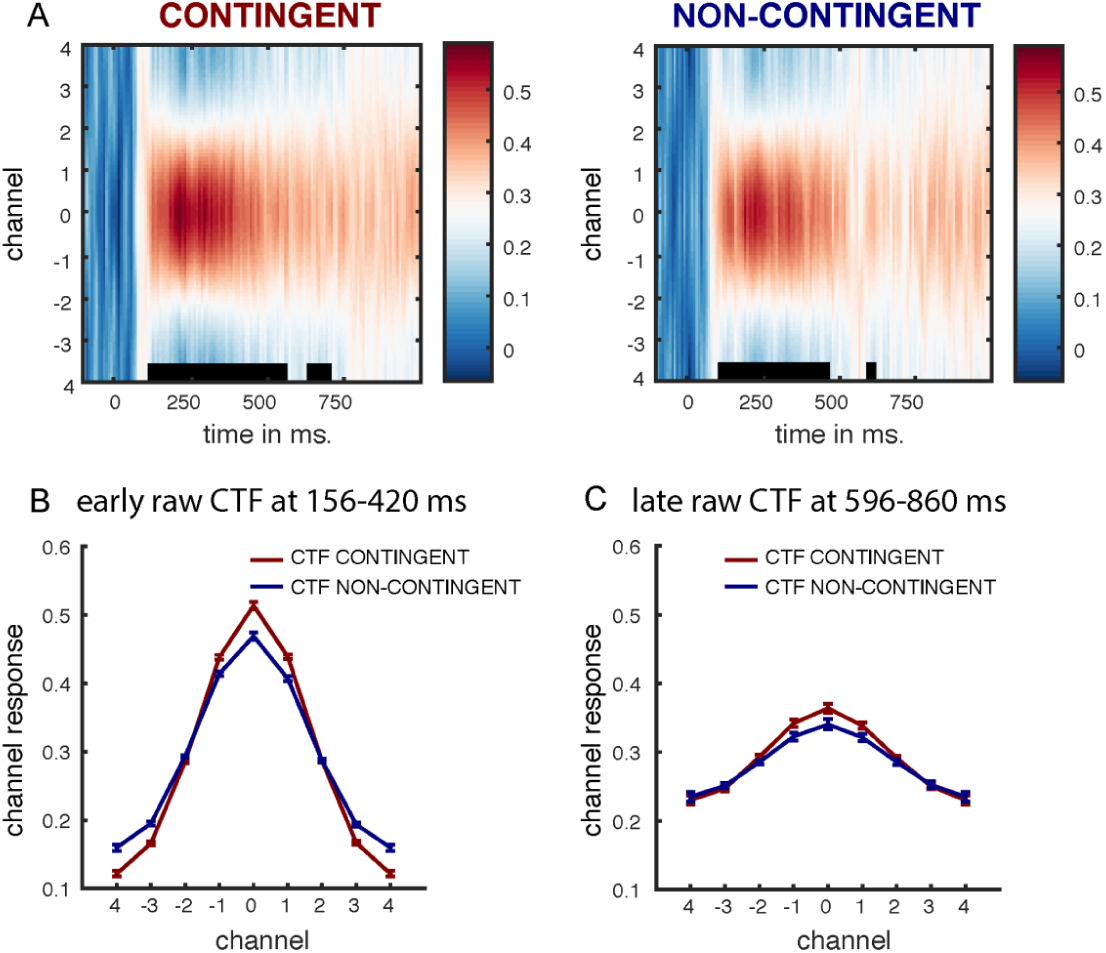
**A**. Development of channel tuning functions from cue onset (tcue = 0) based on raw EEG data. Significance is indicated by black bars on the x-axes. **B.** Contingent and non-contingent slope for the *early* time interval (156-420 ms after cue onset). **C**. Contingent and Non-Contingent slope for the *late* time window (596 – 860 ms after cue onset).

To investigate whether the created forward encoding model describes a continuous relation between cue location and early and late effects of attentional capture, we compared the strength (i.e. the slopes) of contingent and non-contingent cue induced tuning functions for the early and late intervals derived from the BDM analysis (i.e. early: 156-420 and late: 596-860). Figure 5b and 5c show the channel tuning functions for contingent and non-contingent cues for these intervals. We investigated the differences in the CTFs for contingent and non-contingent cues by directly comparing the slopes of each of the obtained functions to each other. In line with the BDM decoding results, the slopes of the evoked CTFs were statistically different for contingent and non-contingent cues in the early interval. A paired samples t-test showed that contingent cues resulted in steeper slopes as compared to non-contingent cues, showing that CTFs were more strongly tuned to the location of contingent compared non-contingent cues (*t*(28) = 5.990, *p* < .001). No such differences were observed in the late interval (*t*(28) = 0.852, *p* = .401), mirroring the results of the raw EEG BDM analyses. Thus, early attentional capture was at modulated by the match between cue color and attentional set.

#### Time-Frequency analysis (FEM)

We subsequently investigated whether a direct and continuous relationship could be established between cue location and the distributed pattern of alpha activity over the scalp. Similar to FEMs based on raw EEG data, and utilizing the same data, channel tuning functions were created separately for contingent and non-contingent cues. As shown in Figure 6A, a strong relationship between cue location and induced alpha-band power was observed yielding robust channel tuning functions for both contingent and non-contingent cues.

**Figure 6.**
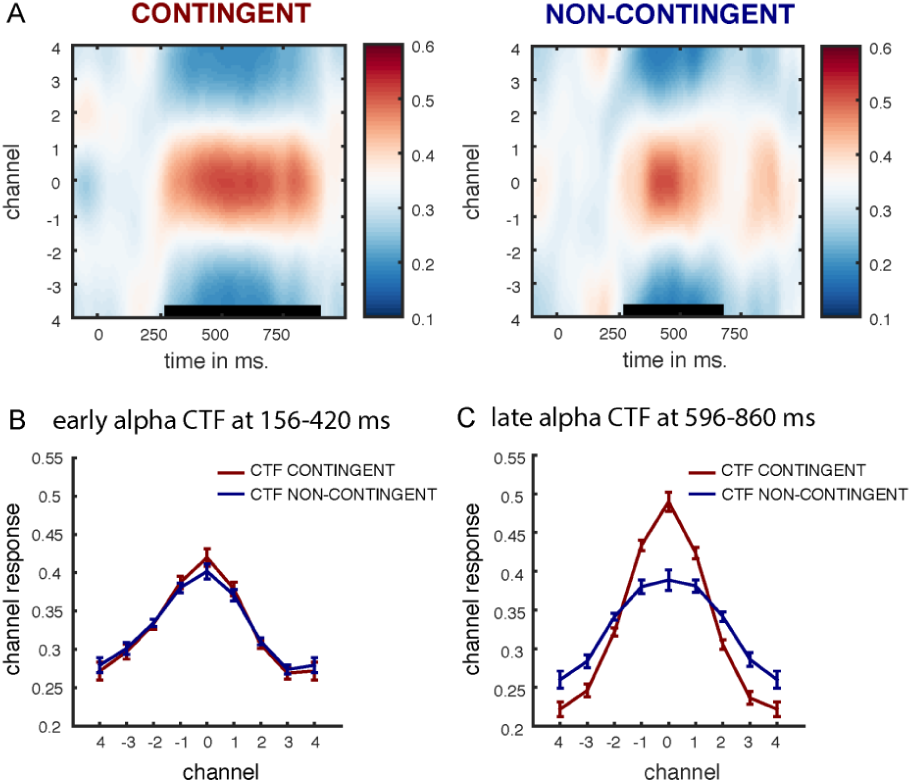
**A**. Development of channel tuning functions from cue onset based on alpha-band power. Significance is indicated by black bars on the x-axes. **B.** Contingent and non-contingent slope for the *early* time interval (156-420 ms post-cue). **C**. Contingent and Non-Contingent slope for the *late* time window (596 – 860 ms post-cue).

Following the results obtained in the BDM analyses and the procedure conducted for the FEM analysis on the raw data, we distinguished between early (156 – 420 ms post-cue) and late (596 – 860 ms post cue) intervals, to equate differences in early effects of attention reflecting automatic capture and late effects of attention that follow initial capture, as encoded in the distributed alpha power. Contrary to the raw EEG encoding results, no difference in slope was observed for the early interval (Figure 6B; *t*(28) = 0.459, *p=* .650), suggesting that alpha-band power did not capture differences in early, automatic attentional modulation of the cues. As expected, a significant slope difference between contingent and non-contingent cues was observed in the late interval (*t*(28) = 2.970, *p =* .006), with contingent cues yielding a steeper slope as compared to non-contingent cues. The results presented in Figure 6b further suggest that there is no difference between the slopes evoked by non-contingent cues in the early compared to the late time interval. However, this lack of an effect is caused by the observation that alpha does not yield strong CTFs early on in the selected time window (> 156 ms), resulting in equally strong CTFs compared to the late time window (the time windows being defined by the BDM analyses and not by the observations presented in Figure 6a).

Analyses of raw EEG and alpha power revealed distinct time courses for the differences between contingent and non-contingent cue conditions. When using the raw EEG signal in the FEM analyses, significant differences between the two cue types can be observed in the early interval (156 – 420 ms after cue onset), but these differences are no longer present during the late interval (596 – 860 after cue onset). The opposite time course was observed with alpha-band power, where reliable differences were observed in the late time interval but not during the early time window. To quantify this inverse pattern, a repeated measure ANOVA was conducted on the individual slope values with contingency (contingent, non-contingent), time interval (early, late) and signal (raw EEG, alpha power). As main effects have already been established, the current results focus solely on the interactions between the different factors. First, the observed early/late reversal for raw EEG/alpha power is supported by a significant three-way interaction between contingency, time interval and signal (*F*(1,28) = 5.791, *p* = .023, *η*_*p*_^2^ = .171). This interaction is illustrated in Figure 7 in which the difference in CTF slope between contingent and non-contingent (contingent – non-contingent) is plotted as a function of time interval and signal. Furthermore, a two-way interaction was observed between signal and time interval, showing that raw EEG yielded steeper slopes in the early time interval as compared to the late time interval, whereas alpha-power resulted in the reversed pattern with steeper slopes in the late, compared to the early time interval (*F*(1,28) = 31.629, *p* <.001, *η*_*p*_^2^ = .530). These results imply that raw EEG and alpha power may track distinct cortical mechanisms related to covert attention. No other interactions were observed.

**Figure 7.**
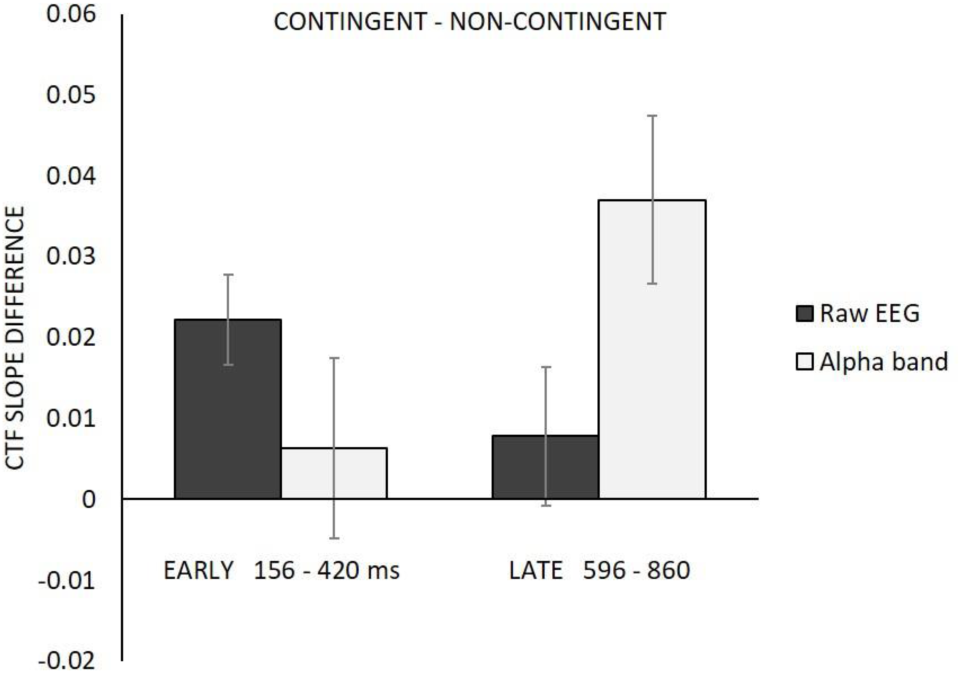
The difference in CTF slopes for contingent and non-contingent cues. A clear interaction between signal type and interval can be observed. Error bars reflect the 95% confidence interval around the mean (Cousineau, 2005; Moray, 2008).

Finally, to ensure that the observed encoding effects are based on a spatially graded profile for each condition (cue location) and to confirm that these effects were not driven by a subset of the cue locations, quadrants or hemifields, we plotted the channel response for each condition separately for early Raw EEG (Figure 8a) and late alpha power (Figure 8b). As can be seen in Figure 8, all locations showed a distinct location-specific CTF in both the raw EEG and the alpha power data, providing further evidence that the observed results in this study indeed reflect location-specific effects of spatial attention.

**Figure 8.**
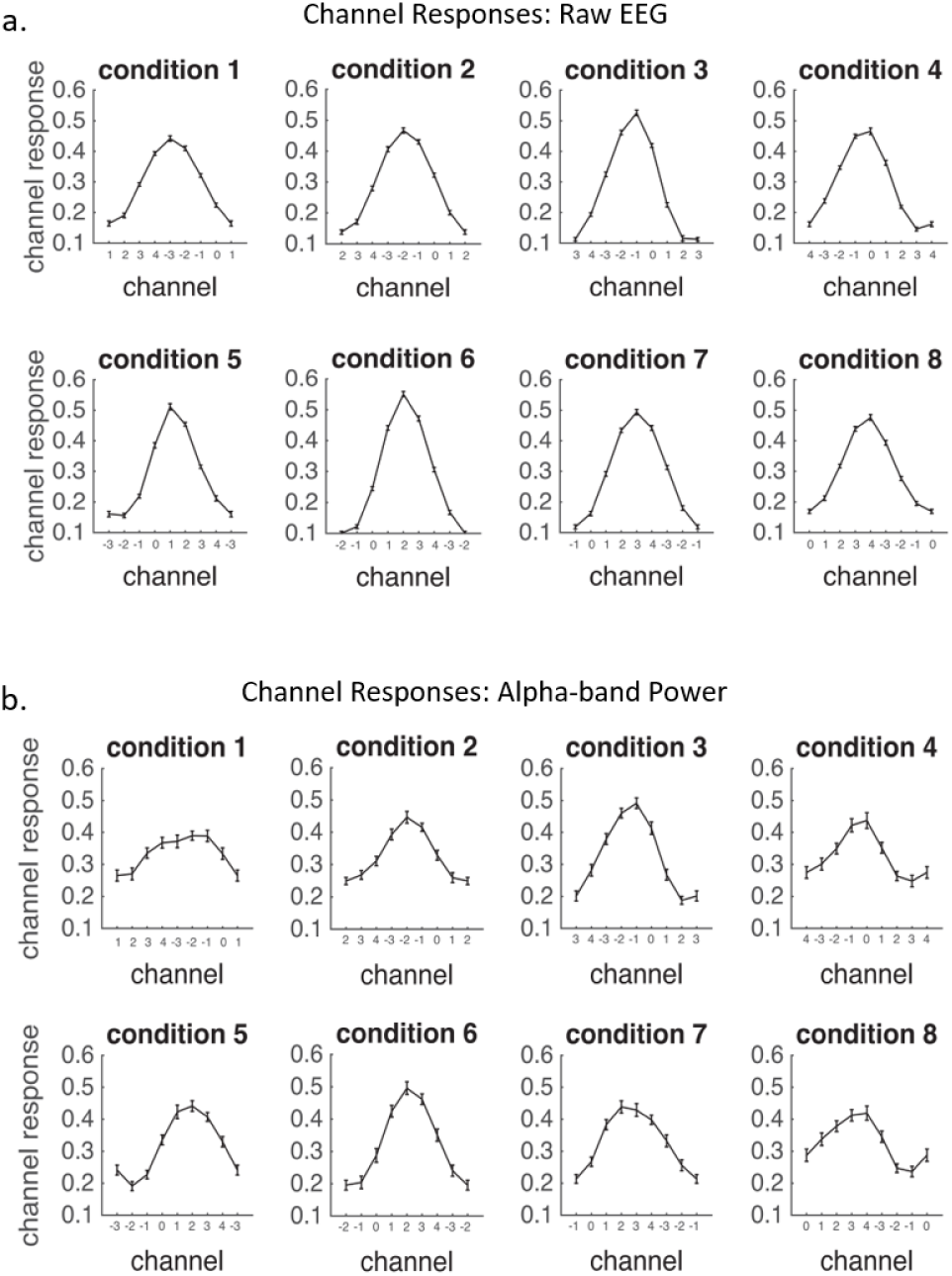
Channel responses for individual conditions showing that the overall CTF is not driven by a subset of lateralized locations, but is based on a full spatial profile in which each location yields a distinct CTF. A) location-based channel responses for early Raw EEG. b) Location-based channel responses for alpha-band power.

## Discussion

The current study aimed to investigate the spatiotemporal properties of attentional capture, with a focus on differentiating between automatic effects of contingent capture and bottom-up capture. To this end, the full spatiotemporal profile of attentional capture was constructed via multivariate analysis of ongoing EEG activity. This approach allowed us to examine attentional allocation in both early and late temporal intervals following the onset of a contingent or non-contingent cue. Modulations of two well-known neurophysiological measures of attention were utilized to establish a direct relationship between attentional processes and neural activity: raw EEG and alpha-power.

Behaviorally, the data mirrored classic results observed in studies that provide support for contingent capture (Egeth & Yantis, 1997; Folk & Remington, 1998, 2006; Folk et al., 1992). Reaction times to the target were influenced by the nature and location of the preceding cue, such that contingent cues exerted a strong influence on reaction times to the target, with reaction times depending on whether they were presented at the location of the target (fast RTs) or elsewhere in the display (slow RTs). This behavioral effect was strongly attenuated for non-contingent cues to the point where non-contingent cues did not produce a reliable RT difference between cues at the target location and cues elsewhere in the display. Critically, however, the results of the EEG analyses clearly indicate that attention was direct towards the non-contingent cue location. As such, the EEG findings provide evidence of attentional capture that was not revealed by behavior alone.

Potential differences in the underlying neural mechanisms responsible for attentional processing of contingent and non-contingent information were assessed in two ways. First, following Fahrenfort and colleagues (2017) and Myers and colleagues (2015), the multivariate distribution of peak values in the raw EEG signal was used to show early and automatic effects of attentional capture. In addition, following Foster and colleagues (2017, 2016), distributed alpha power was used to test the allocation of spatial attention when faced with contingent and non-contingent cues. Backward decoding models were utilized to derive the location of the different cues based on the distributed neural EEG signal, whereas forward encoding models were used to establish a direct and continuous relationship between observed systematic EEG patterns and cue location, separately for each cue type.

By capitalizing on qualitatively different neurophysiological markers in the EEG signal (i.e. the raw EEG data and alpha power), the current study argues against the extreme version of both goal-driven and stimulus-driven models of attentional capture. First, two neural signals known to track the deployment of covert attention revealed that both contingent and non-contingent cues elicited attentional capture. This finding is consistent with the notion that salient signals capture attention independent of attentional control settings (e.g., Theeuwes 2010, 2018). That being said, it is also clear that cues that are contingent with the task set generate stronger and more robust capture even at the very early time points, consistent with the contingent capture hypothesis of Folk et al. (1992). In line with previous work, the multivariate analyses based on the raw EEG (BDM and FEM) showed that the influence of attention emerged shortly after cue onset, but showed its strongest effect in the multivariate EEG patterns around 200-250 ms post cue. This peak time interval matches that of the well-established N2pc component (Eimer, 1996; Luck & Hillyard, 1994) and has repeatedly been linked to general attentional processes, such as identifying and localizing potential target stimuli embedded in an array of non-targets (Eimer, 1996; Hickey, McDonald, & Theeuwes, 2006; Luck & Hillyard, 1994; Mazza, Turatto, & Caramazza, 2009) and more specifically to contingent capture (Grubert et al., 2017; Eimer, 1996; but see Hickey et al., 2006 for a bottom-up account). Nonetheless, differences in decoding between contingent and non-contingent cues can be observed as early as 156 ms after cue onset, preceding the classic N2pc time course which shows its earliest effects around 180 ms after stimulus onset. The most likely explanation for this effect is that the N2pc signal is modulated by an earlier bottom-up signal, evoked by the presentation of the singleton cue stimulus. Therefore, the observed above-chance decoding accuracy is hypothesized to be generated by the same evoked neural activity that is responsible for generation of the N2pc, but precedes it as it is modulated by an earlier bottom up signal. This finding has been substantiated by a recent study by Fahrenfort et al. (2017) in which a forward encoding model (based on raw EEG) was used to describe the continuous relationship between different target locations and systematic fluctuations in neural patterns that arose in the N2pc time window. The current results nicely dovetail those by Fahrenfort and colleagues, by showing a similar continuous relationship between potential target (i.e., the cue) location and systematic fluctuations in the distribution of the raw EEG peaks in the N2pc time window. However, contrary to Fahrenfort et al., the current results not only showed this relationship for contingent trials, but a similar albeit weaker observation was observed for non-contingent cues. This latter finding is most likely caused by properties of the experimental design in which the used cues are the only colored and salient stimuli in the display therefore strongly capturing attention in a bottom-up fashion.

The results of the FEM analysis focused on alpha power converged with those from the BDM analysis. Steeper CTF slopes were observed for contingent as compared to non-contingent cues in the late time window defined in the alpha-based BDM analysis. This observation contrasts with a recent study by Harris and colleagues (Harris, Dux, Jones, & Mattingley, 2017) who showed that alpha-band oscillations were most strongly associated with attentional processes related to contingent cues, but not to non-contingent cues, whereas theta band activity (4-8 Hz) was associated with capture of both cue types, with stronger effects for contingent cues. However, Harris and colleagues investigated alpha and theta oscillatory activity as a function of early attentional capture, focusing on early effects of attention following cue presentation. In our analysis, relatively early (∼250 ms after cue onset) decoding of cue position was observed in alpha activity, but differences in attention to contingent and non-contingent cues were not observed until a later time interval which coincided with the onset of the target stimulus. The observed late effects in the difference between neural activity underlying processing of contingent and non-contingent cues may reflect slower disengagement from contingent relative to non-contingent cue locations. As argued by Theeuwes et al. (Theeuwes, Atchley, & Kramer, 2000) disengagement from the cue may be fast when the cue and the target do not share the same defining properties, while disengagement is slow when cue and target have the same defining features (see also Belopolsky et al., 2010; Fukuda & Vogel, 2009; Grubert et al., 2017). Theeuwes et al., (2000) showed that it takes only about 100 to 150 ms to disengage attention from a cue that does not match the target feature one is searching for.

An alternative and speculative interpretation of the observed late effects in the alpha-based analyses follows recent literature that suggests a link between alpha power and endogenous processes (e.g. David, Kilner, & Friston, 2006; Hosseini, Bell, Wang, & Simpson, 2015). As such, the observed results tentatively suggest a role for slower, voluntary endogenous spatial attentional control that is possibly initiated by early effects of attentional capture. For example, the observed late effects in the difference between neural activity underlying the processing of contingent and non-contingent cues could potentially reflect the effort necessary to disengage from an initially captured, but invalid cue location, with more attentional control being required from a location that contained a contingent cue, as compared to locations containing a non-contingent cue. Perhaps this endogenous signal can be more easily “turned off” when the presented cue has a low target-similarity as in the case of non-contingent cues. That is, observers can rapidly re-allocate spatial attentional resources once it is clear that the attended location does not contain the target. While this is a speculative interpretation of the late alpha based results, the current data suggests that disengaging attention from a target with high target-similarity appears to take more time and effort such that endogenous spatial attention to a contingent may still linger at the cued location when the target is presented (see also, Belopolsky et al., 2010; Grubert et al., 2017).

Nonetheless, while a role of endogenous attention seems appropriate around the moment of target onset, above-chance decoding accuracy based on alpha tuning was observed for both contingent and non-contingent cues starting around 250 ms post cue. An interpretation in terms of voluntary attention seems problematic for this result, as it would not be beneficial to voluntarily attend to the largely uninformative cues (12.5% valid). As such, given the early onset of alpha-based modulations to both contingent and non-contingent cues, it appears that modulations of alpha activity may also reflect processes that are directly linked to early, involuntary bottom-up capture. The discrepancy between the current results and those observed by Harris and colleagues (Harris et al., 2017) is not immediately clear, but may be attributable to different types of analyses used to study spatial attention.

While our data show that contingent cues elicited stronger attentional capture, it may be premature to refer to this as an instance of “goal-driven” attention, because target color was held constant for each observer. Under these conditions, it has been established that selection biases will linger for the selected color even when the current goals of the observer have changed. This lingering effect is sometimes referred to as “selection history” (Awh, Belopolsky, & Theeuwes, 2012), and it describes an interesting class of selection phenomena in which neither physical stimulus salience nor the current attentional goals of the observer can explain the selection bias (see also Theeuwes, 2018). A similar phenomenon can be observed in studies of value-driven capture showing that attention is involuntarily allocated to stimuli that are associated with obtaining a reward, even when the reward is no longer available and the value-signaling stimulus is not the target, ruling out an explanation in terms of top-down effects (Anderson, 2013; Anderson, Laurent, & Yantis, 2011).

In sum, the current study shows that salient stimuli with features that either match or do not match a defining target feature evoke two consecutive but independent attentional signals. First, such stimuli elicit an ‘early’ effect in which attention is captured by all salient stimuli, but with a stronger influence of attention for contingent compared to non-contingent stimuli. This early effect appears to function as input for a ‘late’ attentional mechanism that is potentially endogenous in nature and can be switched off when the attended location does not contain the sought-after target stimulus (or a stimulus closely matching that target). In line with the current results, Hopfinger and West (2006) have suggested that bottom-up and top-down attention can operate concurrently and interactively, by showing distinct and overlapping effects of attention on information processing. The current study elaborates on this finding by showing that attentional capture draws upon multiple attentional mechanisms to shape target selection and identification.

## Footnotes

1 The current method section describes the stimuli and procedure as conducted at the University of Oregon. Minor details, such as the brand of monitor and type of amplifier were different at Bilkent University, and are omitted from the method section. Crucially, stimulus timing and physical properties, as well as the used recording electrodes were identical between testing locations.

2 The procedure for measuring eye movements at Bilkent University was slightly different as it relied on the placement of only one electrode diagonally under the left eye. This procedure proved more than capable in picking up horizontal and vertical eye movements as well as eye blinks.

